# A novel and simple maneuver greatly improves rat liver transplantation

**DOI:** 10.1101/2024.11.13.623355

**Authors:** Yongfeng Chen, Jiabin Zhang, Guoyong Chen, Shaotang Zhou

**Author notes:** Corresponding author: Guoyong Chen, Shaotang Zhou, **Address:** 7 weiwu Road, Jinshui district, zhengzhou.450003, China. Yongfeng Chen and Jiabin Zhang equally contribute to this paper.

## Abstract

**Background:** Orthotopic rat liver transplantation (OLT) is preferentially used to study transplant immunology; it remains challenging due to higher complications associated with 26-minute anhepatic time ceiling, here we safely extended the anhepatic time to improve success of OLT.

**Methods:** Initially we performed OLT including whole graft OLT from inbred Sprague Dawley (SD) rat to SD (whole graft group, control group, n=21) and 30 minute anhepatic time (AHT) group (AHT group, n=11) to master this procedure. For generalization of this maneuver, partial OLT was performed from 50% Lewis allograft to Brown Norway (BN) rat to induce tolerance (half graft group, n=28), Cyclosporine A was injected once daily for 10 days in this group.

**Results:** For whole graft group, 30-day survival rate was 85.5% (18/21), the reasons of death were gas embolism due to the missed suturing in 2 cases, blood loss in 1 case. For AHT group, 30-day survival rate was 72.7% (8/11), one recipient died from respiratory failure intra-operatively; two deaths were biliary leakage in 14 days. There were no differences in survival between whole graft group and AHT group (*p*=0.368). For tolerance group, 30-day survival rate was 82.0% (11/61), and the causes of death were diverse and recorded.

**Conclusion:** The anhepatic time can be extended to 30 minutes simply through the change of clamping the diaphragm, which facilitates LT and improves its success good for the basic research.

## Background

Rat OLT is well-accepted model especially in the basic research of ischemia-reperfusion injury, liver regeneration and immunology since Lee and his colleagues developed in 1973 [1], later in 1979 by Kamada and Calne the introduction of the cuff technique represented a major advance for small animal OLT, simplifying the vascular anastomosis and shortening the portal clamping time [2,3]. Now the two-cuff technique is widely used for OLT in rats, which in this scenario includes 4 or 5 anastomosis of the suprahepatic inferior vena cava (SHVC), the portal vein, infrahepatic inferior vena cava (IHVC), the bile duct and/or the liver artery, any delay of the venous anastomosis will extend or lead to longer clamping time of PV that is taken as longer AHT confering a negative impact on outcomes [4-6]. Short AHT as a surgical skill has been prioritized to minimize as soon as possibly by clinicians and microsurgeons who perform liver transplantation. Clinically it ranged from 37 to 321 minutes, it was reported that over 100-minute AHT was associated with a higher incidence of graft dysfunction [7]. Experimentally, the AHT ceiling of rat OLT is 26 minutes in literature, cardiac arrest occurred in the recipient unexpectedly when AHT was more than 26 minutes [8]. Higher mortality rate during long AHT necessitates extending AHT; short AHT plays a critical role of the success of LT and improves survival greatly. After two-cuff technique introduction, rapid reconnection of SHVC is the most difficult and critical point for OLT. Here we first introduce a method to extend AHT to 30 minutes while simplifying this complicated procedure to improve higher success rate.

## Methods

For our project, rat OLT was performed to induce immunological tolerance through stem cells and liver regeneration (detailed protocol out of scope here). In literature, we observed that clamping the diaphragm led to abnormal breathing and rat death (supplementary 1), and that the change of clamping the diaphragm simplified this microsurgical procedure and improved rat survival. Whole OLT was performed from close SD to SD with different AHT, and partial OLT was performed with Lewis to BN as the acute rejection model whereas this technique was generalized and applied to our project (Table 1).

**Table 1.** Data for recipients of 3 groups. AHT, 30- minute anhepatic time, * *p*=0.368 for survivals between whole graft and AHT group.

|  | Anhepatic time | 30-day survival rate | Death reasons |
| --- | --- | --- | --- |
| whole graft (n=21) | ≤20 min | 85.5% (18/21) | missed suture(2),bleed(1) |
| AHT group ( n=11) | 30 min | 72.7%* (8/11) | respiratory failure(1), biliary leakage(2) |
| half graft (n=61) | ≤20 min | 82.0%(11/61) | wound(6),bile leak(4),infection(1),unknown(2) |

### Animal

Male rats including SD, Lewis and BN (weighing 200-400g) were used as donors and recipients purchased from Laboratory Animal Technology Corporation. The animals were housed with controlled temperature and light, freely access to a standard chow diet and water; they were fasted 12 hours prior to the operation. All experiments were approved by the ethics committee (HNTCMDW-20190304) of Henan Provincial People’s Hospital and conducted in compliance with the standards for animal use and care set by ARRIVE guidelines and the Institutional Animal Care Committee.

### Surgical procedure

Anesthesia was applied with isoflurane inhalation with air to the donors and with oxygen to the recipients. In the donor procedure, under microscope we preferentially select a transverse incision to enter the abdomen of the rat. Firstly through the aorta, a researcher flushed the donor liver with heparinized normal saline (250 u/ml) and then through PV reflushed with lactated Ringer’s solution. The graft was explanted out and stored in a dish full of lactated Ringer’s solution. At the back table, PV and IHVC were respectively everted and secured on cuffs with different inner and outer diameters. For 50% graft, the caudate lobes, the left lateral one and the left portion of the median lobe were removed. Cold storage time was within 3 hours in all cases. In the recipient, to showcase this simple surgical maneuver, we selected the midline incision to open the abdominal cavity [9]. The proper liver artery was ligated proximally, and the accessory liver artery was ligated and cut, a blunt separation behind the liver SHVC was made to create a tunnel. The recipient PV and IVC were blocked respectively with micro-vascular clamps, and isoflurane was immediately decreased to 0.2 volume %. A mosquito forceps was placed through the tunnel on the part (1/3) of diaphragm ring (left side) to occlude SHVC (Figure 1), SHVC was anastomosed with 8-0 polypropylene running suture (Figure 2), when this anastomosis was completed, the forceps was displaced with a vascular bulldog on the real SHVC while the diaphragm ring was de-clamped (Figure 3), the anhepatic time was generally less than 20 minutes. PV was reconnected with the cuff; blood flow was restored once the clamp on the PV was released. IHVC reconnection was made as was for PV. For AHT group, the clamp on the PV was released until 30 minutes. The gastroduodenal artery was proximally ligated and an opening was made on the common hepatic artery into which the stent in the donor hepatic artery was inserted and secured (Figure 4, 5), bile duct continuity was made when a tube in the donor bile duct was inserted into the recipient bile duct [9]. The abdomen was closed with two layers and the animals were kept in a cage under an infrared light. 10% glucose solution and purified water were both supplied for 3 first days, and later regular food and tap water were offered. Sodium ceftizoxime (100mg/kg) was injected subcutaneously once a day, 4 days in total. For half graft group, cyclosporine A was subcutaneously injected once daily for 14 days and ceased afterwards. Sacrifice was made on deep anesthesia with isoflurane for euthanasia.

**Figure 1.**
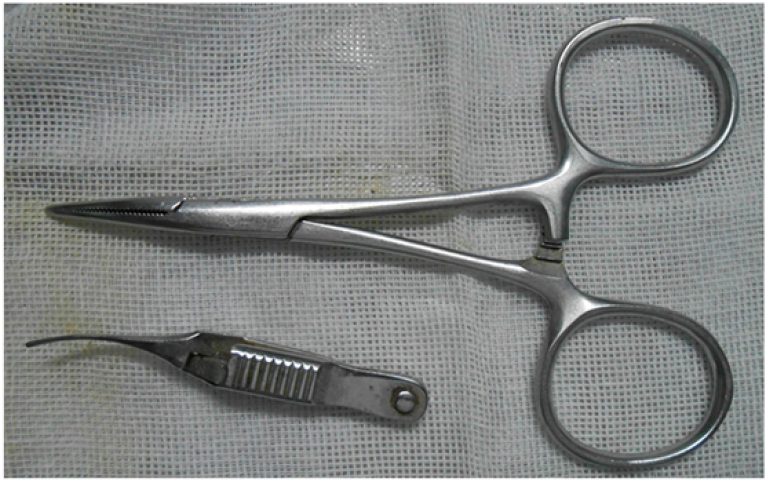
The mosquito forceps and the bulldog for occlusion of SHVC.

**Figure 2.**
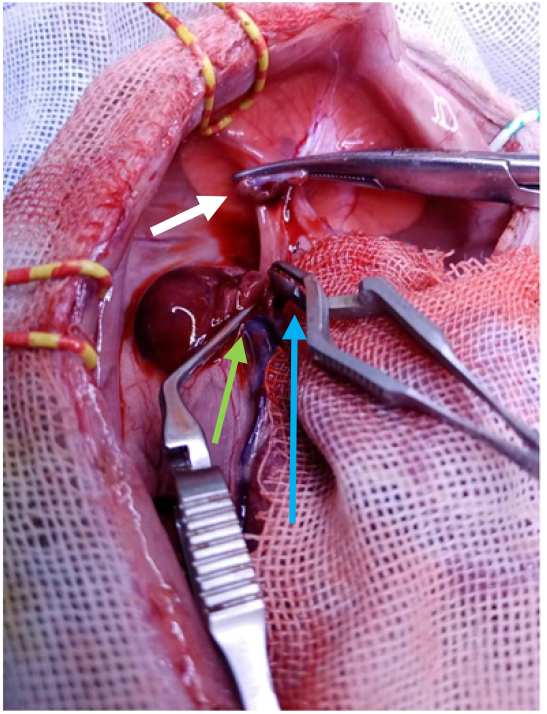
Clamping the partial diaphragmatic ring with the mosquito forceps while the native liver was removed. White arrow for SHVC, green one for IHVC, blue one for PV.

**Figure 3.**
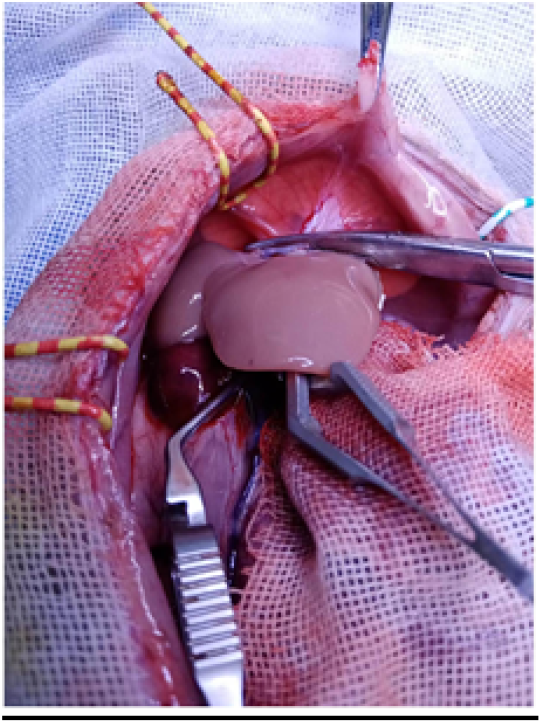
The donor liver graft was placed orthotopically.

**Figure 4.**
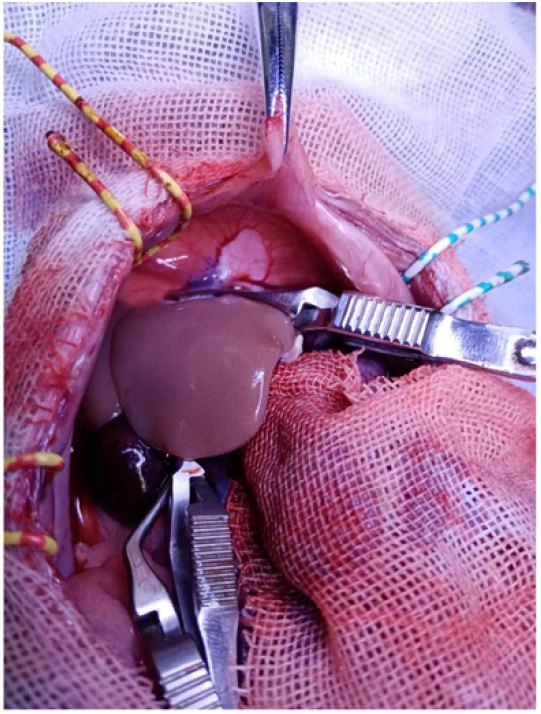
The diaphragmatic ring was declamped and the real SHVC was clamped with the bulldog.

**Figure 5.**
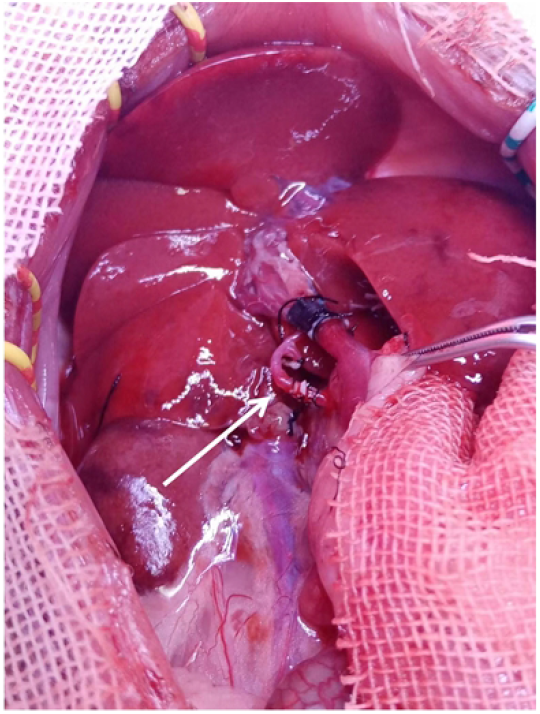
The view of LT after circulation is restored. The artery was reconnected (arrow).

### Statistical analysis

The cumulative survival rates of different groups was evaluated with the Kaplan-Meier Curve, the analysis was made with SPSS 22.0 software (IBM Corp, Armonk, NY, USA), and *P*<0.05 was considered significant

## Results

In the beginning, a microsurgeon selected the transverse incision for OLT, he displaced the clamping forceps to occlude SHVC that was hand-sutured and became skillful over time, later he selected the midline incision for less trauma. For the whole graft, 30-day survival rate was 85.5% (18/21), the reasons of death were gas embolism due to missed suturing during the anastomosis in 2 cases, blood loss in 1 case. For AHT group, 30-day survival rate 72.7% (8/11), one recipient died from respiratory failure intra-operatively; two deaths were biliary leakage in 14 days. There were no differences in survival between whole graft group and AHT group (*p*=0.368) (supplementary 1). For half graft group, 30-day survival rate was 82.0% (11/61) (Figure 6), the reasons of death were euthanasia due to self-biting in 6 cases respectively on day11,12,13,17,18,31; biliary leakage occurred to 4 cases on day 22,26,45,46, infection to one case on day 13, unknown reasons to 2 case on day 13,43 post-LT (supplementary 2, 3). Histological examination revealed almost normal structures of liver without fibrosis, ductopenia, thickened wall in the liver arterioles and venules in half graft group (H&E staining not shown).

**Figure 6.**
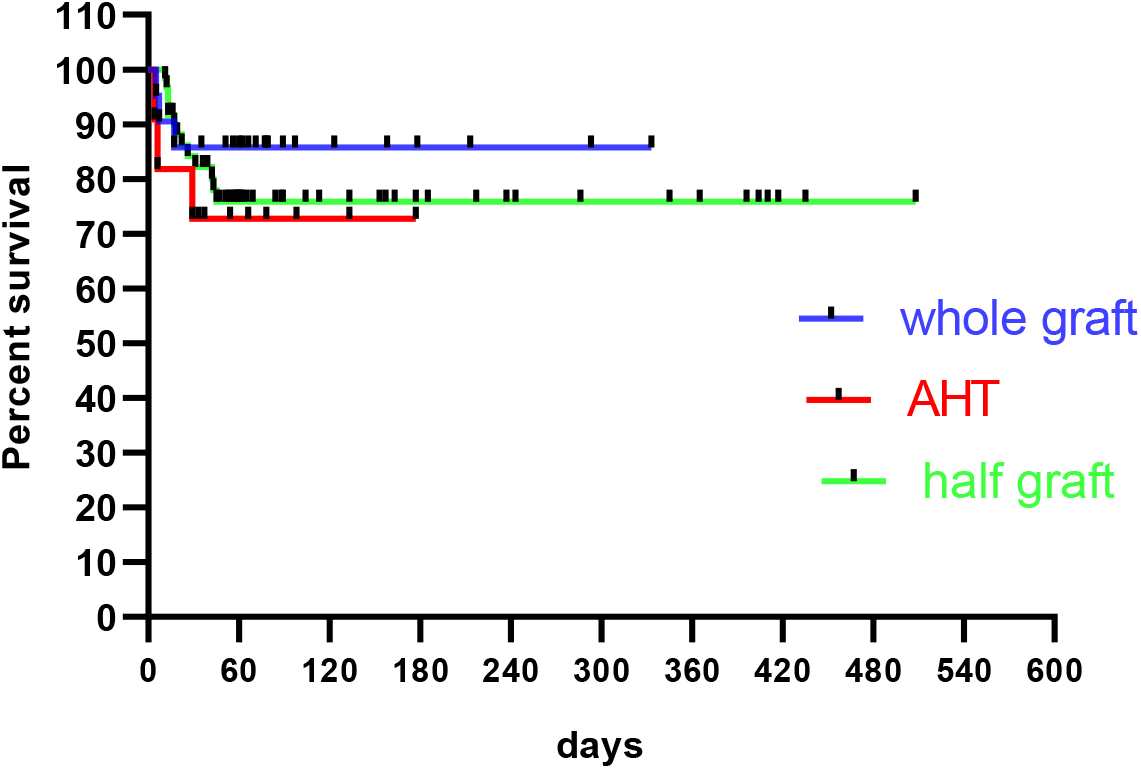
Survivals for different groups.

## Discussion

Kamada et al introduced the cuff method for OLT to greatly benefit OLT for basic research [1-6], however it is still complicated especially when it is performed by a microsurgeon without an assistant surgeon. Suturing SHVC by hand is the first and tough procedure to reconnect vessels for OLT due to short SHVC. While SHVC is blocked without ventilator aid, it necessitates to clamp the diaphragmatic ring that maintains respiratory function for better exposure, once clamped and retracted downward (caudalad) to perform anatomosis, it dramatically affects respiratory and the rat starts to move, you had to discontinue suturing, some inexperienced surgeons might be much nervous to add anesthesia, consequently it has a big impact on respiratory or the rat will die (supplementary 1), especially under plain and simple mask anesthesia inhalation (ether etc), this is the main reason that respiratory insufficiency or cardiac arrest occurred to lead to higher mortality during AHT [8]. Generally, clamping partial diaphragmatic ring is prerequisite and good to better expose SHVC for suturing directly circulation is temporarily halted and it will affect respiratory, the recipient rat will surely move a few seconds, you had to wait for its motionlessness. When the reconnection of SHVC is completed, the diaphragmatic ring is de-clamped immediately and the real SHVC is clamped with a fine bulldog to avoid continuous respiratory disturbance or avoid air embolism, subsequently the surgeon takes time to reconnect PV and IHVC. For our report, it is safely extended to 30 minutes or more whereas survival rates were not significantly different, in the time-efficient manner (30 min), the different anhepatic time has little impact on recipient survival. Over time, you will become more efficient and the success rate is approaching or beyond 90%.

Alternatively the cuff method may comprise the reconstruction of SHVC [4,10-15], Cuff method efficiently shortens SHVC anastomosis, but it is not universally applicable and easier due to short SHVC in length. Magnetic ring is a cuff method in nature and precludes future MRI examination [13, 15].

Rapid anastomosis of vessels and short AHT reflect microsurgical skills in OLT. Some researchers took some measures to extend AHT for outcome improvements of OLT. Prostaglandin and its analogue were used to extend AHT and improve survival [16,17]. Liu et al reported that clamping the supra-celiac aorta one minute can effectively improve rat OLT by increasing the tolerable time of AHT [6], this maneuver does not surpass the AHT ceiling. Our maneuver is surgically easy and effectively extends AHT to 30 minutes (supplementary 2, 3). A few references were reviewed to reveal that the diaphragm was clamped much more [18, 19], we followed that procedure which led to irregular breathing (supplementary 1). In literature almost no attention has been paid to how to clamp SHVC or the diaphragm upon the reconnection of SHVC, our report is the first description of clamping the diaphragm. In human liver transplantation, a strong blocking forceps is placed on the diaphragm and secured stably, it will have little impact on respiratory because of availability of assisted breathing via ventilator.

As above-mentioned, the complication of self-biting is related to the transverse incision due to greater trauma, its occurrence is avoided when the midline one is selected; biliary leakage is related to the shorter donor bile duct and its reduction was recorded on longer bile duct.

For tolerance induction in our report, it is ongoing research of our project and the detailed protocol is out of scope here.

## Conclusion

OLT can be safely performed with extension to 30 minute AHT simply through the change of clamping the diaphragm; this procedure facilitates LT and improve its success rate greatly.

## Abbreviations

OLT: orthotopic liver transplantation
AHT: anhepatic time
SD: Sprague Dawley
BN: Brown Norway
PV: portal vein
SHVC: supra-hepatic vena cava
IHVC: infra-hepatic vena cava

## Declarations

### Ethics approval and consent to participate

All experiments were approved by the ethics committee of Henan Provincial People’s Hospital (HNTCMDW-20190304) and conducted in compliance with the standards for animal use and care set by ARRIVE guidelines and the Institutional Animal Care Committee of Henan Provincial People’s Hospital. All methods were performed in accordance with the relevant guidelines and regulations.

### Consent for publication

**N/A**

### Availability of data and materials

All data and materials are available on request.

## Author contributions

YF. Chen conceived the study and revised the draft. JB. Zhang analyzed data and revised the draft. ST Zhou performed OLT, wrote the draft, designed and finalized the study. GY. Chen funded and discussed the study. All authors have read and approved the manuscript.

## Acknowledgements

N/A

## Supplementary Legends

**Video 1. T**he diaphragm was clamped regularly (like references) and the reconnection of the SHVC was finished, the irregular respiratory occurred and the rats died when the diaphragm was de-clamped.

**Video 2**. The rat respiratory was normal when clamping was performed like my study.

**Video 3**. View after the liver artery was reconnected with a stent.

